# Structural Basis for Membrane Recruitment of ATG16L1 by WIPI2 in Autophagy

**DOI:** 10.1101/2021.05.14.444175

**Authors:** Lisa M. Strong, Chunmei Chang, C. Alexander Boecker, Thomas G. Flower, Cosmo Z. Buffalo, Xuefeng Ren, Andrea K. H. Stavoe, Erika L. F. Holzbaur, James H. Hurley

## Abstract

Autophagy is a cellular process that degrades cytoplasmic cargo by engulfing it in a double membrane vesicle, known as the autophagosome, and delivering it to the lysosome. The ATG12–5-16L1 complex is responsible for conjugating members of the ubiquitin-like ATG8 protein family to phosphatidylethanolamine in the growing autophagosomal membrane, known as the phagophore. ATG12–5-16L1 is recruited to the phagophore by a subset of the phosphatidylinositol 3-phosphate-binding seven bladed β-propeller WIPI proteins. We determined the crystal structure of WIPI2d in complex with the WIPI2 interacting region (W2IR) of ATG16L1 comprising residues 207-230 at 1.85 Å resolution. The structure shows that the ATG16L1 W2IR adopts an alpha helical conformation and binds in an electropositive and hydrophobic groove between WIPI2 β-propeller blades 2 and 3. Mutation of residues at the interface reduces or blocks the recruitment of ATG12–5-16L1 and the conjugation of the ATG8 protein LC3B to synthetic membranes. Interface mutants show a decrease in starvation-induced autophagy. Comparisons across the four human WIPIs suggest that WIPI1 and 2 belong to a W2IR-binding subclass responsible for localizing ATG12–5-16L1 and driving ATG8 lipidation, whilst WIPI3 and 4 belong to a second W34IR-binding subclass responsible for localizing ATG2, and so directing lipid supply to the nascent phagophore. The structure provides a framework for understanding the regulatory node connecting two central events in autophagy initiation, the action of the autophagic PI 3-kinase complex on the one hand, and ATG8 lipidation on the other.

## INTRODUCTION

Macroautophagy (hereafter autophagy) maintains cellular homeostasis by sequestering unneeded or harmful cytoplasmic material in double membrane vesicles known as autophagosomes (Morishita and Mizushima, 2019). Mature autophagosomes fuse with lysosomes, so degrading their contents. Starvation-induced autophagy is thought to target bulk cytosol, while various forms of selective autophagy target damaged mitochondria and other organelles, invading bacteria, protein aggregates, and many other intracellular materials (Anding and Baehrecke, 2017; Gomes and Dikic, 2014). Defects in autophagy are associated with increased vulnerability to pathogens, aging, and neurodegenerative diseases (Levine and Kroemer, 2019). Defects in the autophagy of mitochondria (“mitophagy”) downstream of Parkin and PINK1 are associated with hereditary early onset Parkinson’s Disease (Pickrell and Youle, 2015; Stavoe and Holzbaur, 2019).

The many varieties of bulk and selective autophagy all rely on a handful of shared core components, which include the class III phosphatidylinositol 3-kinase complex I (PI3KC3-C1); the ubiquitin-like ATG8 family (LC3A-C, GABARAP, and GABARAPL1-2 in mammals); the proteins ATG7, ATG3, and ATG12–5-16L1 responsible for conjugating ATG8s to phosphatidylethanolamine (PE); and the WD-repeat protein interacting with phosphoinositide (WIPI family) (Chang et al., 2021a; Mizushima et al., 2011). PI3KC3-C1 is targeted to sites of autophagy initiation by its ATG14 subunit, where it phosphorylates phosphatidylinositol (PI) at the third position in the inositol ring to generate PI(3)P (Itakura et al., 2008; Obara et al., 2006; Sun et al., 2008). ATG8 proteins are attached to the membrane lipid phosphatidylethanolamine (PE) in a process that is closely analogous to the conjugation of ubiquitin to its target proteins (Ichimura et al., 2000). In brief, ATG4 cleaves ATG8 to expose the C-terminal glycine, the ubiquitin E1-like ATG7 then activates ATG8 for transfer to the ubiquitin E2-like ATG3, and the ATG12–5-16L1 complex scaffolds the ATG8 transfer from ATG3 to the headgroup of PE (Klionsky and Schulman, 2014). The function of ATG12–5-16L1 is analogous to that of ubiquitin E3 ligases, and we therefore refer to this complex here as “E3”. This process is often referred to as LC3 lipidation, after LC3, the founding member of the ATG8 family in mammals (Kabeya et al., 2000). In mammals, ATG8 conjugation to membranes is important for multiple steps in autophagy, and is particularly critical for autophagosome-lysosome fusion (Nguyen et al., 2016; Tsuboyama et al., 2016).

The two critical steps in autophagy initiation, PI 3-phosphorylation and LC3 lipidation, are connected to one another via a direct interaction between a subset of the PI(3)P-binding WIPI proteins and ATG16L1 (Dooley et al., 2014). The human WIPI1-4 proteins comprise a subset of the seven bladed β-propeller protein binding to phosphoinositides (PROPPINs) (Dove et al., 2004). PROPPINs bind to PI(3)P and PI(3,5)P_2_ headgroups through a conserved FRRG motif (Dove et al., 2004; Gaugel et al., 2012) and bind tightly, but reversibly, to membranes using a hydrophobic loop in blade 6 that inserts into the membrane (Baskaran et al., 2012; Krick et al., 2012; Watanabe et al., 2012). WIPI2 is expressed as six known isoforms, which appear to have overlapping functions (Proikas-Cezanne et al., 2015). WIPI2b in particular has been shown to have a central role in bulk and selective autophagy initiation in cells (Dooley et al., 2014; Polson et al., 2010), and WIPI2d potently activated LC3 lipidation in an *in vitro* giant unilamellar vesicle (GUV) reconstituted system (Fracchiolla et al., 2020).

Despite the centrality of the WIPI2:ATG16L1 interaction to mammalian autophagy initiation, only a predictive model (Dooley et al., 2014), but no experimentally determined structure has been available. Here, we report the crystal structure of WIPI2d: ATG16L1 (207-230) complex at a 1.85 Å resolution. WIPI2d point mutations in the interface disrupted ATG16L1 binding, reduced the ability of WIPI2 to recruit ATG12–5-16L1 and promote LC3 lipidation on GUVs, and reduced starvation-induced autophagy in cells.

## RESULTS

### Structure determination of WIPI2d:ATG16L1-W2IR

In order to generate a crystallizable form of WIPI2d, the flexible hydrophobic loop in blade 6 and the putatively disordered C-terminal region were deleted (Fig. 1A). The deletion construct removes the only regions whose sequence diverges between WIPI2b and WIPI2d, thus the construct represents a WIPI2b/d consensus. A peptide corresponding to the WIPI2-interacting region (“W2IR”) comprising residues 207-230 of ATG16L1 (Dooley et al., 2014) was synthesized. The crystal structure of the WIPI2d: ATG16L1 complex was determined at 1.85Å (Fig. 1B, C) by molecular replacement using the structure of *Kluveromyces lactis* Hsv2 (Baskaran et al., 2012) (PDB: 4EXV) as a search model. ATG16L1 was modelled *de novo* into the density (Fig. 1D). The asymmetric unit contains two copies of the WIPI2d: ATG16L1 W2IR complex. One WIPI2d monomer is bound to one ATG16L1 peptide, the two copies align with a Ca root-mean-square deviation (RMSD) of 0.3 Å. Statistics of crystallographic data collection and structure refinement are provided in Supplementary Table 1. As expected on the basis of the Hsv2 (Baskaran et al., 2012; Krick et al., 2012; Watanabe et al., 2012) and WIPI3 (Ren et al., 2020) structures, WIPI2d folds into a seven blade β-propeller, with each blade containing four anti-parallel β-strands. The propeller is ~50 Å wide and ~30 Å tall (Fig. 1B, C). The FRRG motif that enables WIPI2d binding to phosphoinositides is distal to the ATG16L1 binding site.

**Figure 1:**
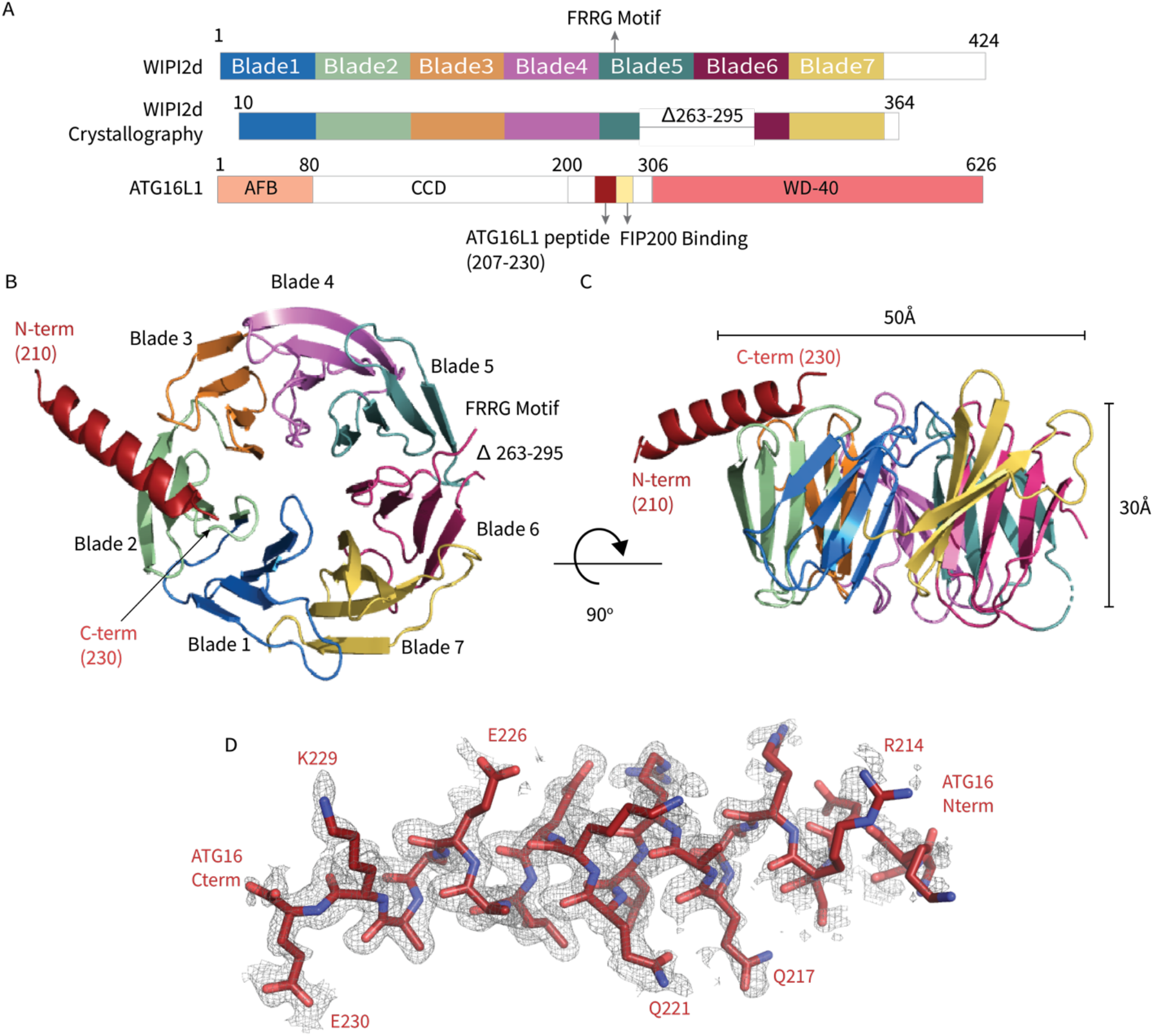
WIPI2d: ATG16L1 W2IR Structure. Structure of WIPI2d bound to ATG16L1 W2IR. A) Annotated WIPI2d and ATG16L1 domain schematics. WIPI2d construct for crystallography is shown and W2IR from ATG16L1. B-C) The ribbon diagram of the WIPI2d complex with ATG16L1 W2IR from the B) bottom and C) side views. Each blade is colored in accordance with A. D) Composite omit map of ATG16L1 W2IR. Modelled ATG16L1 is shown as red carton and the composite omit 2mFo-DFc map contoured at 1σ is shown in grey.

### Analysis of WIPI2d W2IR: ATG16L1 Interface

The ATG16L1 W2IR nestles between blades 2 and 3 of WIPI2d, burying ~550 Å^2^ of solvent-accessible surface area. Blades 2 and 3 are identical in all six WIPI2 isoforms, thus, we expect that conclusions concerning the ATG16L1 binding mode drawn here will pertain to all WIPI2 isoforms. The WIPI2d binding site for the ATG16L1 W2IR consists of a single deep groove with a mixed electropositive and hydrophobic character (Fig. 2A, C). Hydrophobic side chains of Leu 64, Phe 65, Leu 69, Val 83, Ile 92, Cys 93, Ile 124, and Met 127 on WIPI2d contribute to the hydrophobic surface of the groove. The surfaces of Leu 220 and Leu 224 of the ATG16L1 W2IR are buried in this interface (Fig. 2C, D). The side-chains of WIPI2d His 85, Lys 88, Arg 108, and Lys 128 contribute to the electropositive character of the groove. The acidic side chains of Glu 226 and Glu 230 of ATG16L1 interact with the electropositive patch on WIPI2 (Fig. 2E). The presence of WIPI2d Arg 108 and Arg 125, and ATG16L1 Glu 230 in the binding site was correctly predicted by the modeling efforts of Tooze and colleagues (Dooley et al., 2014). The nature of their interactions can now be defined on the basis of the crystal structure of the complex. Gln 217 of ATG16L1 forms a hydrogen bond with Lys 128 of WIPI2d at the N terminus of the W2IR and WIPI2d, respectively. The C-terminus of the ATG16L1 W2IR, Glu 230 forms a salt bridge with Arg 108 of ATG16L1. Arg125 makes a water-mediated bridge to the W2IR peptide backbone in one of the two complexes in the asymmetric unit. Ser 66, Ser 67, and Ser 68 contribute additional polar interactions. The backbone of ATG16L1 near Ala 227 and Ala 228 forms a hydrogen bond with the backbone of WIPI2d between residues Ser 68 and Leu 69. This backbone binding favorably buries residues Leu 64, Phe 65, and Ser 67 within WIPI2d.

**Figure 2:**
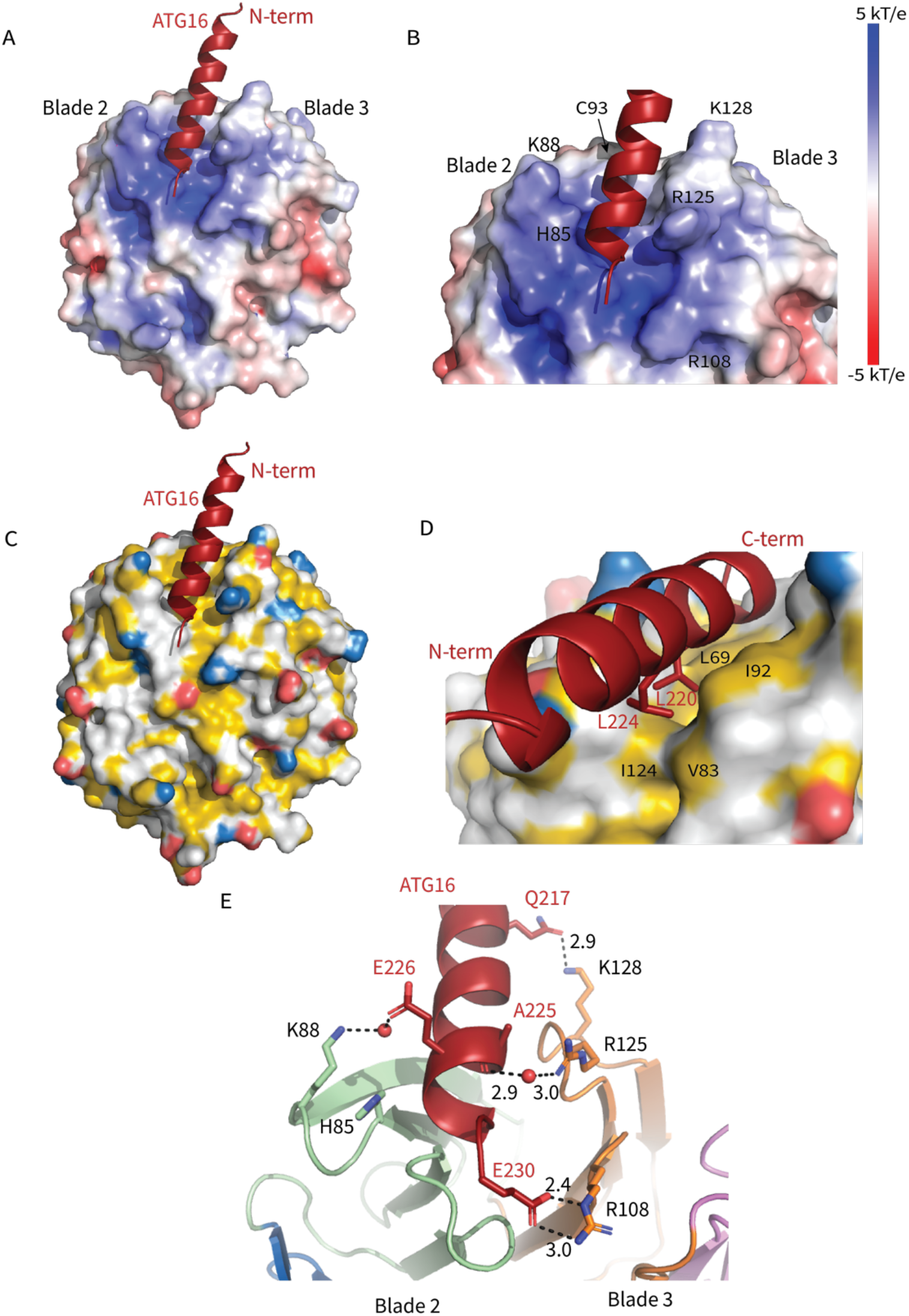
Interactions at the interface. Analysis of the Interface. A) Overall electrostatic surface and B) closer view of electrostatic surface with ATG16 W2IR shown as a cartoon and key residues labelled. C) Overall hydrophobic surface of WIPI2d and D) closer view of the hydrophobic interface with key residues labelled where yellow represents hydrophobic regions. E) A cartoon and stick representation of hydrogen bonds between ATG16 and WIPI2d shown as black dotted lines with distances noted and key residues shown as sticks.

### Roles of WIPI2 interfacial residues

To evaluate the role of specific residues at the interface, we introduced single site mutations into WIPI2d to disrupt binding. H85E, K88E, and C93E were designed to perturb the electropositive WIPI2d surface on blade 2 (Fig. 2B, 3A, B). L69E and I92E were designed to disrupt the hydrophobic groove for hydrophobic packing of ATG16L1 (Fig. 2D, 3A, B). K128E and R108E were chosen to abolish the interactions with Gln 217 and Glu 230 in ATG16L1, respectively (Fig. 2E, 3A, B). R125E was designed to disrupt the bridging interaction to Lys88 (Dooley et al., 2014). Both R108E and R125E were previously been shown to reduce binding within the cellular context, thus these two mutants also served to confirm that our *in vitro* binding experiments can replicate the findings of previously reported immunoprecipitations (Dooley et al., 2014). To investigate the complex formation of these mutants, we purified these mutants and performed a coprecipitation assay using immobilized GST-ATG16L1 W2IR (Fig. 3C, D). It was observed that L69E and C93E were prone to aggregation and were therefore not characterized further. All other mutants expressed at near identical levels as wild-type, were purified at equivalent yields, and so presumed not to have grossly perturbed structures and stabilities. H85E, K88E, and I92E completely abolished binding to ATG16L1 while R108E and R125E retained weak binding to ATG16L1 (Fig. 3D). Interestingly, K128E binds with similar affinity to WT WIPI2d (Fig. 3D). Lys128 is positioned within a flexible β-loop (Fig. 3A) near the location of three disordered Arg residues in the N-terminal part of the ATG16L1 W2IR preceding Gln 217. The resulting charge repulsion might offset the contribution of the W2IR Gln 217 hydrogen bond. The presence of these apparent negative interactions suggests that the affinity of the wild-type complex has evolved to be moderate to facilitate the dissolution of the complex during the course of autophagosome maturation.

**Figure 3:**
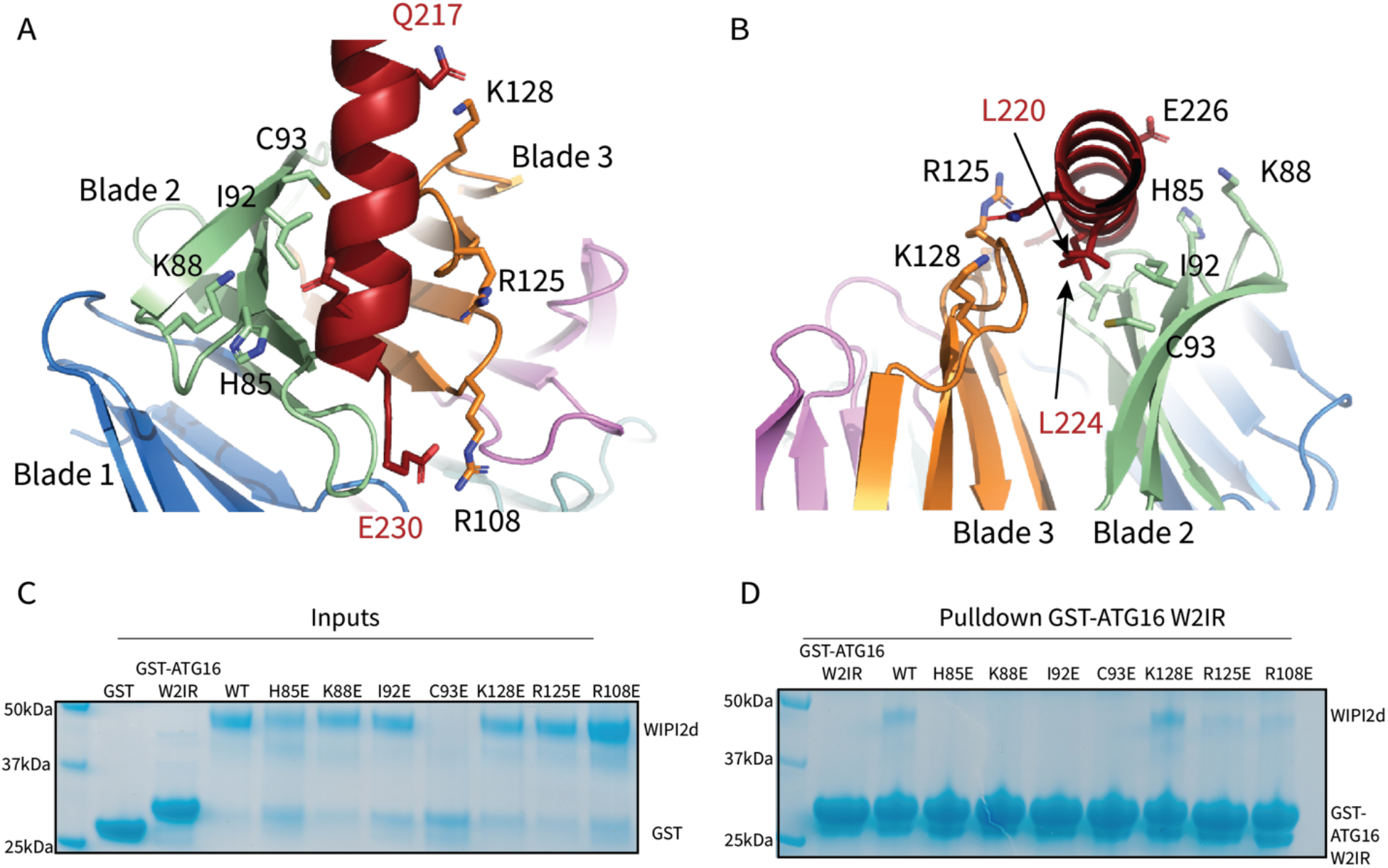
WIPI2d Interfacial mutants decrease ATG16L1 binding. Key interacting residues shown as sticks in cartoon representation of WIPI2d: ATG16L1 interface shown from A) the WIPI2d face or B) down the ATG16L1 helix. C) Pull-down assays of mutant WIPI2d constructs and wild type with GST-ATG16L1 W2IR. GSH resin was used to pull down GST-ATG16L1 W2IR from purified protein mixture. The pull-down results were visualized by SDS-PAGE and Coomassie blue staining.

### The WIPI2d: ATG16L1 W2IR interface is required for LC3 lipidation *in vitro*

We next assessed the ability of WIPI2d mutants to activate E3 membrane recruitment and LC3 lipidation in a microscopy-based GUV assay (Chang et al., 2021b; Fracchiolla et al., 2020). In the presence of WIPI2d WT and the LC3 conjugation machinery (ATG7, ATG3, The ATG12–5-16L, and a mCherry-LC3B construct corresponding to the ATG4-processed form) (Fig. 4A), PI3KC3-C1 robustly triggered membrane recruitment of the E3-GFP complex and activated mCherry-LC3B lipidation (Fig. 4B, C). Consistent with expectation, mutation of the previously characterized ATG16L1 binding sites R108E and R125E significantly reduced E3 membrane binding and LC3 lipidation (Fig. 4B, C). The mutants H85E and I92E almost completely abolished E3 membrane binding and LC3 lipidation (Fig. 4B, C). The K88E mutant also had an obvious defect in E3 recruitment and LC3 lipidation (Fig. 4B, C). All of these observations are consistent with the loss of binding noted in the GST pull-down experiments. Consistent with the positive pull-down result, the K128E mutant fully retained the ability to recruit E3 to GUV membrane and activate subsequent LC3 lipidation (Fig. 4). These data confirm that the ATG16L1 binding interface on WIPI2d is responsible for the E3 recruitment and activation that promotes LC3 membrane conjugation.

**Figure 4:**
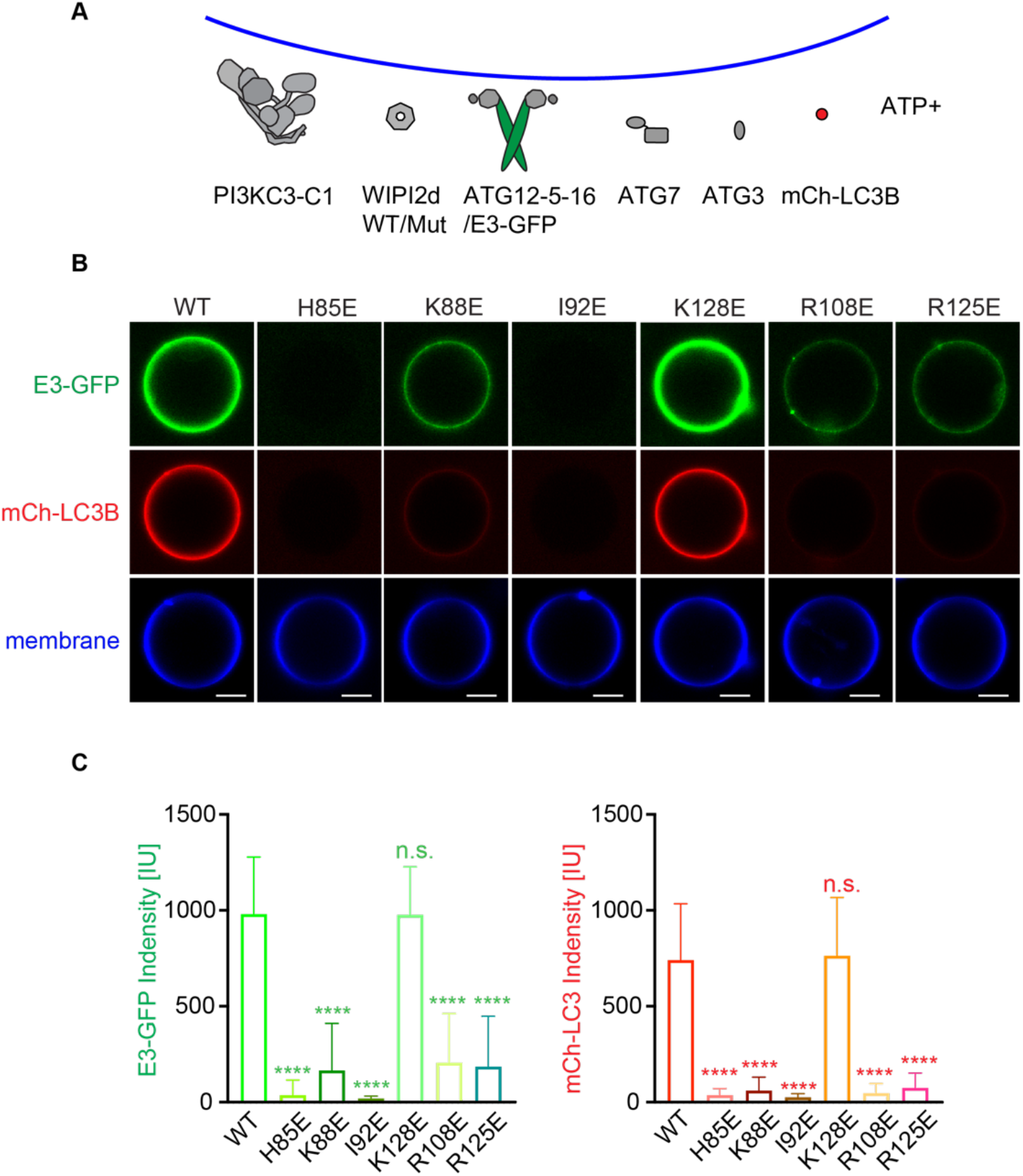
WIPI2d mutants disrupt E3 recruitment and LC3 lipidation on GUVs. A) The schematic drawing illustrates the reaction setting. Colors indicate fluorescent protein fused components. Components in gray are not labeled but are present in the reaction. B) Representative confocal images of GUVs showing E3 membrane binding and LC3B lipidation. PI3KC3-C1, WIPI2d WT or mutant, E3-GFP, ATG7, ATG3, mCherry-LC3B and ATP/Mn^2+^ were incubated with GUVs (64.8% DOPC: 20% DOPE: 5% DOPS: 10% POPI: 0.2% Atto647 DOPE) at room temperature. Images taken at 30 min were shown. Scale bars, 10 μm. C) Quantification of relative intensities of E3-GFP and mCherry-LC3B on GUV membranes in (A) (means ± SDs are shown; N = 40). p≥0.5: (ns); 0.01<p<0.05: (*); 0.001 <p<0.01: (**); p<0.001 (***); p<0.0001 (****).

### Mutations that disrupt the WIPI2: ATG16L1 W2IR interface impair starvation-induced autophagy

Together, our structural observations and in vitro reconstitution experiments predict that mutations that disrupt the electrostatic interface between WIPI2 and ATG16L1 will disrupt autophagosome formation in vivo. To test this hypothesis, we engineered H85E, K88E, I92E, C93E, and K128E mutations into WIPI2B. We expressed Halo-tagged mutant constructs in parallel to WT WIPI2B in mouse embryonic fibroblasts (MEFs) depleted for endogenous WIPI2 by siRNA (Fig. 5A,B). In parallel, we also expressed the R108E mutation previously demonstrated by Dooley et al. (Dooley et al., 2014) to disrupt the WIPI2: ATG16L1 interaction. All mutants tested expressed at levels similar to WT (Fig. 5C). Autophagy was induced via a 2-hour incubation of the MEFs in starvation media (EBSS) in the presence of 100 nM Bafilomycin A (BafA). WIPI2 puncta formation was scored, with the lowest levels of puncta formation observed in cells expressing I92E or C93E (Fig. 5D). As expected, none of the mutations abrogated the recruitment of WIPI2, but the lower numbers of WIPI2 puncta seen in cells expressing I92E or C93E may reflect a more transient localization of these mutant proteins to the omegasome (Fig. 5A). Next, we examined the formation of LC3-positive autophagosomes (Fig. 5A,E). Here, we found that every mutation except I92E induced a significant inhibition of autophagosome formation, with the most pronounced deficits seen upon expression of the C93E and R108E mutations; the H85E, K88E, and K128E mutants all showed similar deficits in autophagosome formation (Fig. 5E). Together with the structural and in vitro data described above, these cellular assays support the model that the electrostatic interface between WIPI2 and ATG16L1 mediates efficient autophagosome formation. However, the observation that autophagy is inhibited but not abrogated upon expression of these mutants suggests that multiple combinatorial interactions facilitate the assembly and function of the complex autophagosome biogenesis machinery in cells, reflecting a relatively robust mechanism for autophagosome formation.

**Figure 5:**
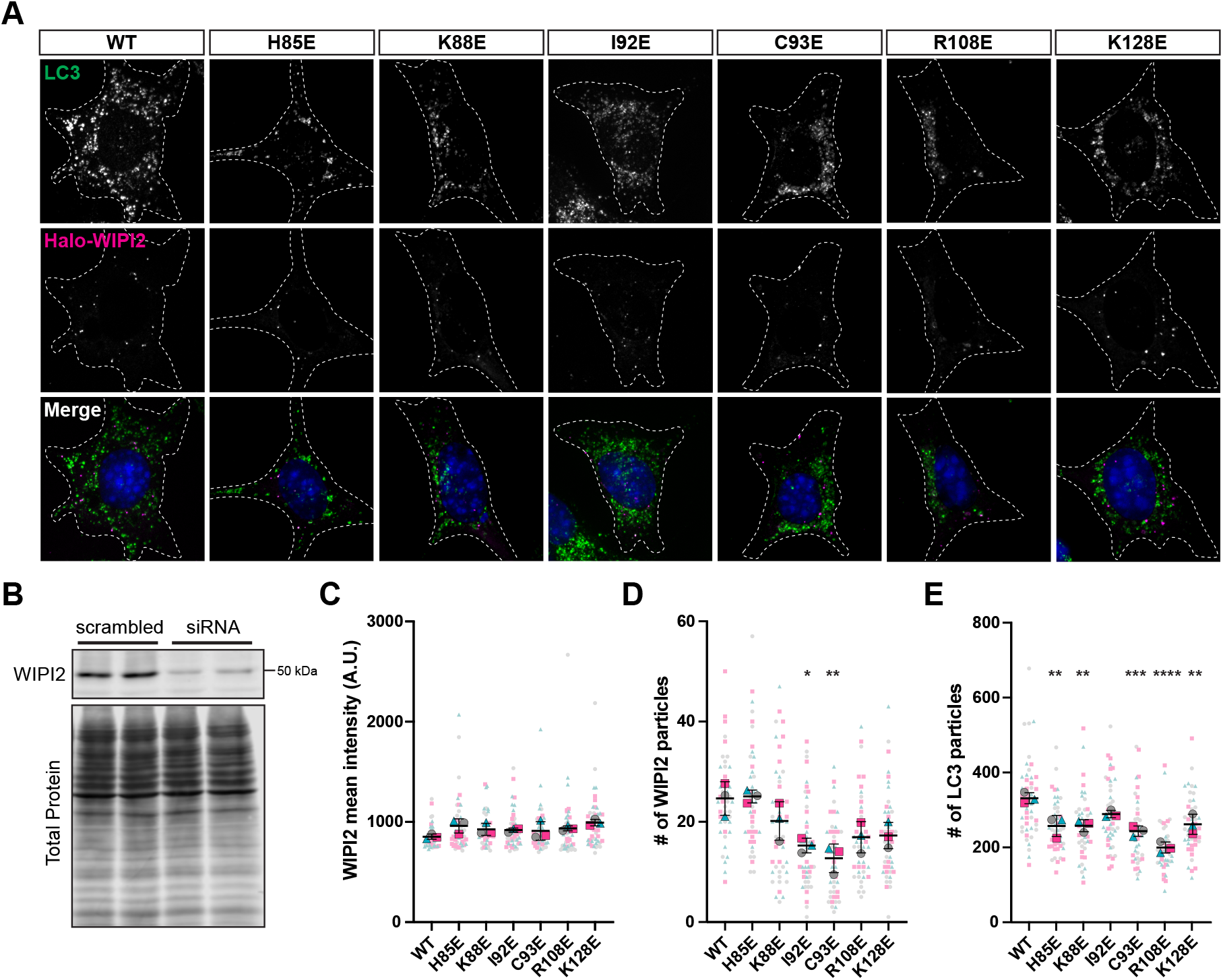
Altering the electrostatic interface of WIPI2 impairs starvation-induced autophagy in MEFs. A) Representative maximal projections of LC3 staining and Halo-WIPI2 signal in MEFs following induction of autophagy via 2-hour starvation in EBSS in the presence of 100 nM BafA. Endogenous WIPI2 was depleted by siRNA and cells were transfected with Halo-WIPI2 WT, H85E, K88E, I92E, C93E, R108E, or K128E. B) Immunoblot of MEF lysates treated with indicated siRNA, collected 48 hours after siRNA transfection. C-E) Quantification of C) the mean fluorescence intensity of Halo-WIPI2, D) Halo-WIPI2 particle number, and E) LC3 particle number in EBSS + BafA starved MEFs, depleted of endogenous WIPI2 and expressing siRNA-resistant WT or mutant Halo-WIPI2. Independent experimental replicates are color-coded, with individual data points in light colors and averages of three independent repeats in bold colors (mean ± SEM; n = 3 independent experiments; *, p < 0.05; **, p < 0.01; ***, p < 0.001; ****, p < 0.0001 between WIPI2 point mutants and WIPI2 WT by one-way ANOVA with Tukey’s multiple comparisons test).

### *In vitro* reconstitution of WIPI2 membrane recruitment

The finding that certain WIPI2 mutants had reduced membrane recruitment led us to examine whether WIPI2 recruitment to GUV membranes was perturbed by the W2IR binding site mutations. A minimal system including PI3KC3-C1 and E3 was used to explore the possibility that even in the presence of PI(3)P, E3 binding might contribute to WIPI2 recruitment. K88E, R108E, and R125E decreased WIPI2 recruitment to a significant extent (Fig. 6A, B), while other mutants did not.

**Figure 6:**
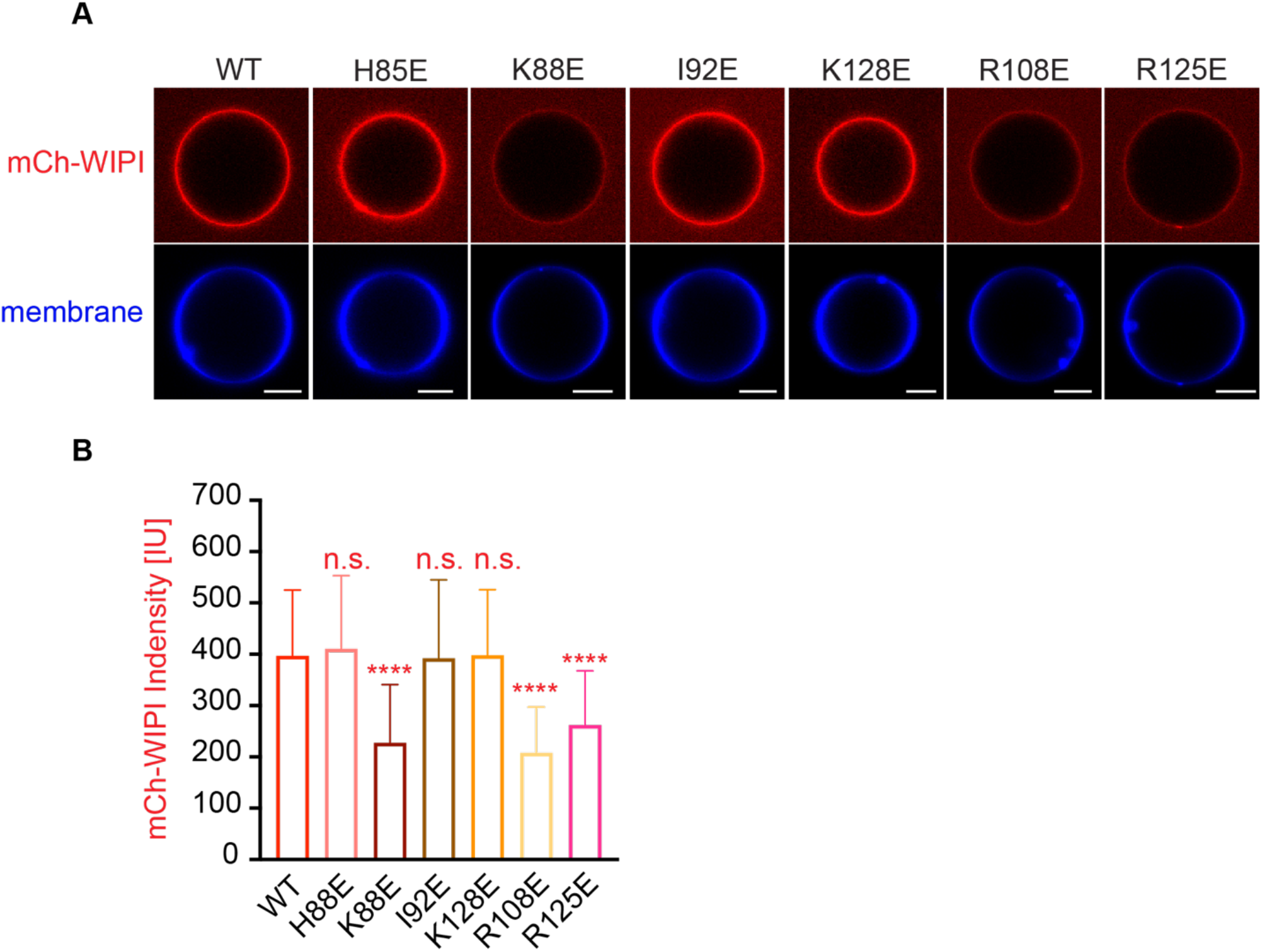
WIPI2d Mutants Membrane Affinity. A) Representative confocal images of GUVs showing membrane binding of mCherry-WIPI2d. PI3KC3-C1, mCherry-WIPI2d WT or mutant, E3-GFP were incubated with GUVs (64.8% DOPC: 20% DOPE: 5% DOPS: 10% POPI: 0.2% Atto647 DOPE) at room temperature. Images taken at 30 min were shown. Scale bars, 10 μm. B) Quantification of relative intensities of mCherry-WIPI2d on GUV membranes in (A) (means ± SDs are shown; N = 40). p≥0.5: (ns); 0.01<p<0.05: (*); 0.001<p<0.01: (**); p<0.001 (***); p<0.0001 (****).

### Comparison across the WIPI protein family

The structure reported here was based on a construct corresponding to a consensus of the WIPI2b/d sequences for blades 1-7, since the C-terminal extension, the only region of divergence between the two proteins was deleted. These are the two WIPI2 isoforms that have been previously shown to bind ATG16L1 in immunoprecipitations from cells (Dooley et al., 2014). While the remaining WIPI isoforms diverge from the 2b/d consensus in blade 1, their sequences are identical in the blades 2 and 3 involved in ATG16L1 binding site. To the extent that these other isoforms were reported not to bind ATG16L1, these differences cannot be inherent in the W2IR binding groove itself, but rather must reflect other differences in the cellular context and modifications.

The only other human WIPI for which a structure is known is that of WIPI3 (Liang et al., 2019; Ren et al., 2020). WIPI3 interacts with the lipid transporter ATG2A (Ren et al., 2020) via what is believed to be a conserved binding site also present in WIPI4. WIPI4 is responsible for recruiting the phospholipid conduit ATG2A to sites of phagophore initiation, where it promotes tethering of the nascent phagophore to the ER membrane source (Chowdhury et al., 2018; Zheng et al., 2017). The structure of WIPI3 is superimposable on that of WIPI2d with a Cα r.m.s.d. of 1.2 Å (Fig. 7A, B).

**Figure 7:**
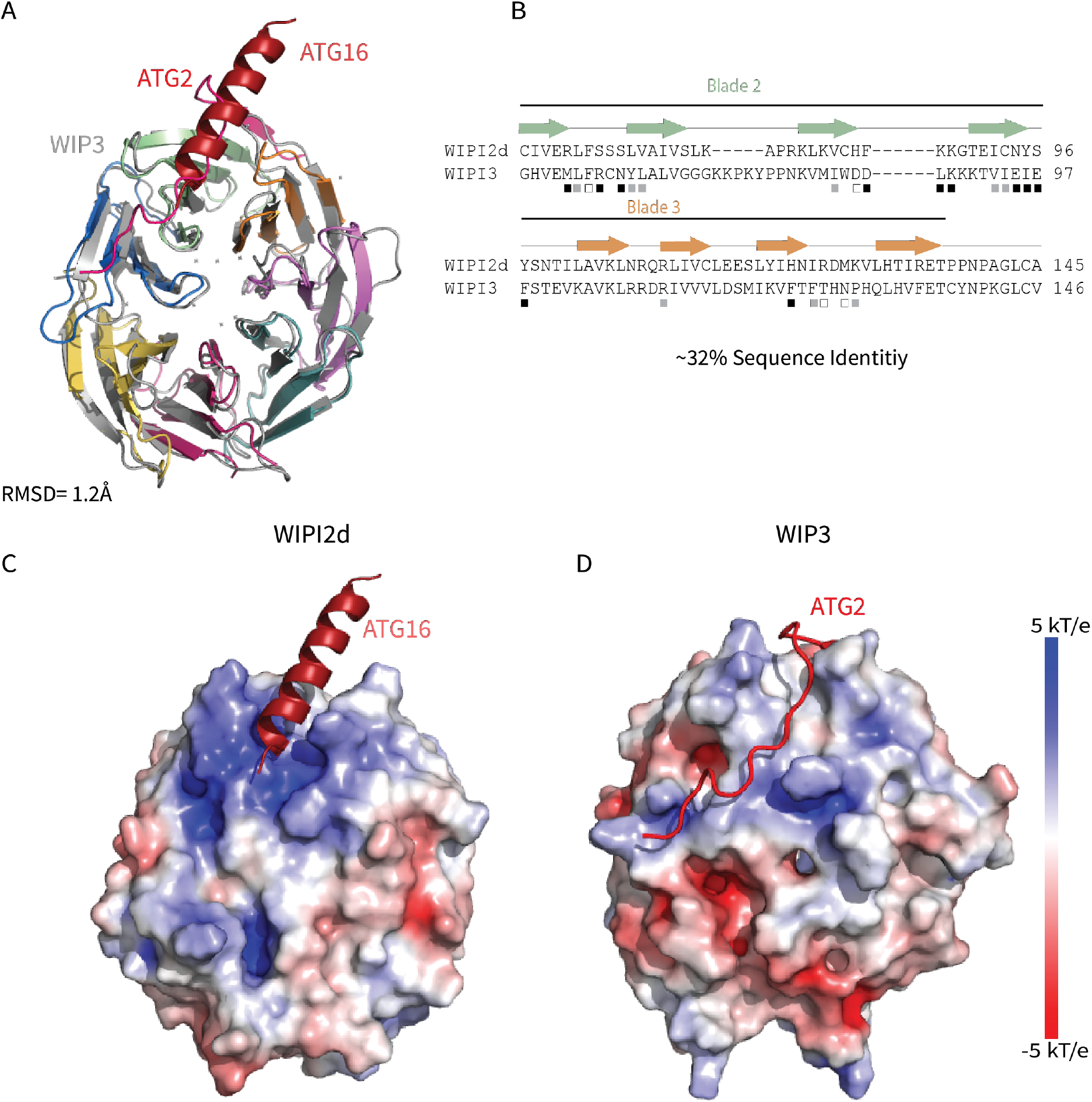
Comparing WIPI2d and WIPI3 structures and binding modes. Comparison of WIPI2d and WIPI3. Alignment of WIPI2d and WIPI3 A) structure and B) sequence based on structures with W2IR residues denoted with white squares, W34IR with black, and from both with grey. Electrostatic surface comparison of C) WIPI2d and D) WIPI3.

## DISCUSSION

WIPI2 is the linchpin of the circuit that connects two of the key reactions in autophagy initiation, the synthesis of PI(3)P by PI3KC3-C1, and LC3 lipidation by ATG12–5-16L1. The WIPI2-ATG16L1 interaction is essential for starvation-induced bulk autophagy and xenophagy (Dooley et al., 2014), and for efficient LC3 lipidation in a reconstituted system with physiologically reasonable nanomolar concentrations of autophagy core complexes (Fracchiolla et al., 2020). From the perspective of therapeutic restoration of autophagic function in aging and neurodegeneration, ectopic expression of WIPI2b restores a normal rate of autophagosome biogenesis in aged neurons (Stavoe et al., 2019). Here, we report the high resolution crystal structure of human WIPI2 and show how its unique electropositive and hydrophobic groove between blades 2 and 3 binds to the ATG16L1 W2IR.

The functional relevance of the groove residues was investigated by *in vitro* LC3 lipidation assays and by LC3 puncta formation in starvation induced autophagy. All but one of the binding site mutants, K128E, reduced in vitro binding as judged by pull down assays of purified proteins. WIPI2 activation of LC3 lipidation of GUV membranes by ATG2-5-16L1 precisely mirrored the results of the pull-down assays, with K128E again being the only mutant exhibiting no reduction. *In vivo* LC3 puncta formation was also reduced by most of the mutants, although the pattern did not follow the same rank order as the *in vitro* results. We interpret these data as confirmation that the W2IR binding site is important for LC3 lipidation *in vivo*, but that the many additional autophagy initiation components present in cells still modulate the effects in subtle ways. In a simple linear paradigm of autophagy initiation, PI(3)P generated by PI3KC3-C1 recruits WIPI2, which in turn recruits E3 to catalyze LC3 lipidation. In this model, mutations that perturb the E3 binding of WIPI2 would not be expected to alter the recruitment of WIPI2 itself. However, at least one other upstream component, FIP200 (Fujita et al., 2013; Gammoh et al., 2013; Nishimura et al., 2013), contributes to E3 recruitment, and ATG16L1 has inherent membrane binding of its own (Lystad et al., 2019). Thus the presence of E3 can stabilize WIPI2 on membranes in cells, a finding bolstered by our observation of the same effect *in vitro*.

Remarkably, the binding site for ATG2A is between blades 2 and 3 of WIPI3, the same two blades involved in binding ATG16L1 by WIPI2 (Fig. 7C, D). Despite the overall close similarity in the folds of the two WIPIs, the detailed structure of the blade 2-3 groove is quite divergent, explaining why WIPI3 does not bind ATG16L1, and WIPI2 does not bind to ATG2A. The Val- and Pro-rich ATG2A sequence that binds to WIPI3 in an extended conformation (Ren et al., 2020), and presumably WIPI4, is completely different in character from the Leu- and Glu-rich helical W2IR of ATG16L1. We propose the term WIPI3/4 interacting region (W34IR) for the ATG2A binding motif to contrast it with the distinct W2IR of ATG16L1. The ATG2A binding groove of WIPI3 is electrostatically neutral, as compared to the electropositive groove in WIPI2. *A* subset of the essential W2IR binding residues of WIPI2 (Fig. 7, white squares) are altered in WIPI3. For example, the critical His 85 of WIPI2 is replaced by Asp in WIPI3. Expanding the analysis to WIPI1 and 4, the main features of the WIPI2 ATG16L1 binding groove are preserved in WIPI1 but not WIPI4 (Fig. 8). Conversely, the ATG2A binding groove of WIPI3 is preserved in WIPI4 but not WIPI1 (Fig. 8). Thus, the structural findings are consistent with the concept that the four human WIPIs can be subclassified into two groups (Polson *et al*., 2010): an ATG16L1-binding WIPI1/2 group and an ATG2A-binding WIPI3/4 group.

**Figure 8:**
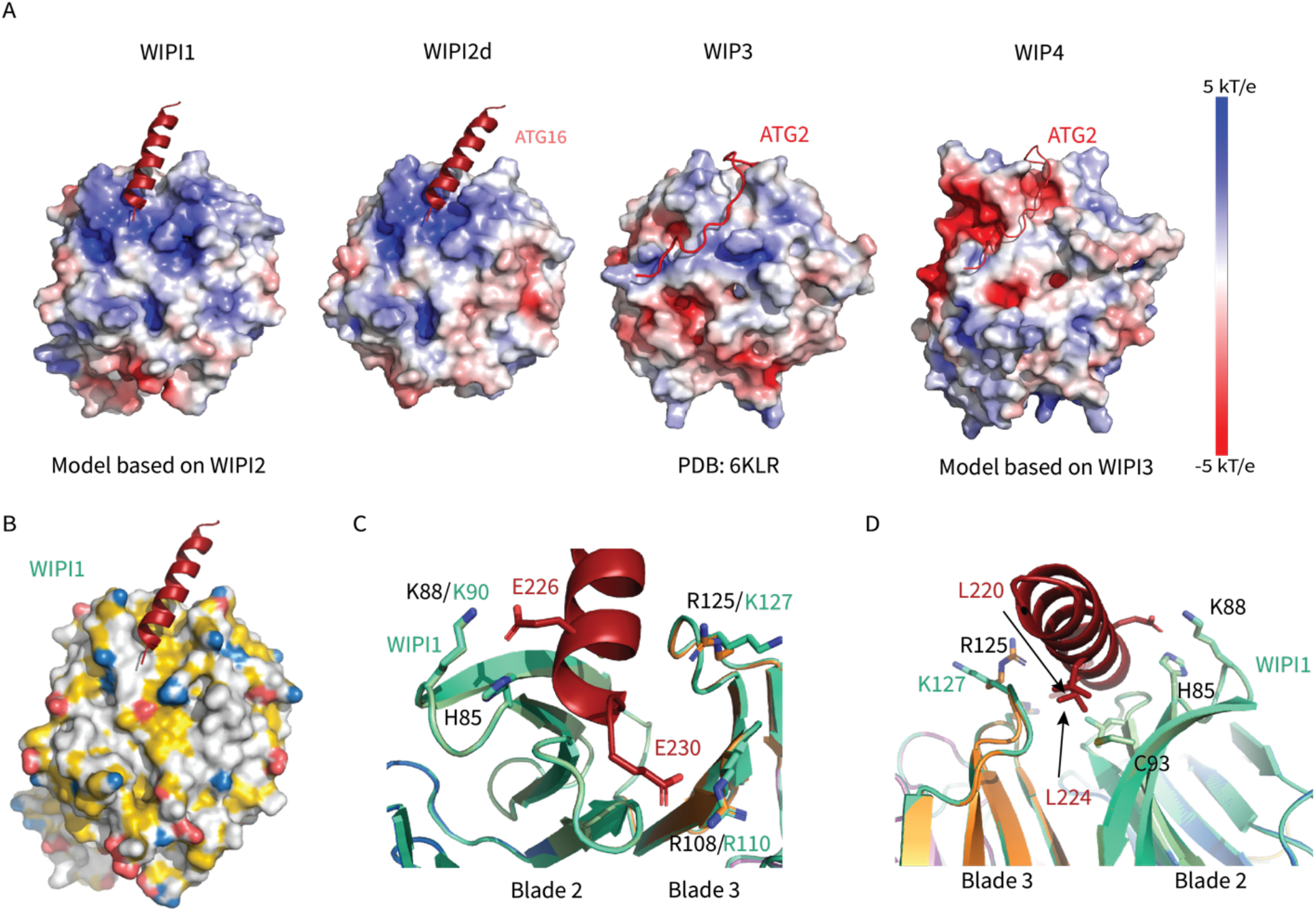
WIPI1-4 comparison. Comparison of electrostatic surface potential of A) WIPI1-4. B) Hydrophobic surface of WIPI1 with predicted ATG16L1 W2IR shown as cartoon. C & D) Alignment of WIPI2d crystal structure and WIPI1 homology structure with WIPI1 shown as light green and key residues labelled in the same color as structure.

Whilst WIPI-based recruitment of ATG16L1 is critical for autophagy, a number of other factors are also involved. FIP200 can recruit ATG16L1 to sites of phagophore initiation (Fujita et al., 2013; Gammoh et al., 2013; Nishimura et al., 2013) via the central region of ATG16L1 that centers on residues 239-246 (Fujita et al., 2013) and so adjoins with the WIPI2 binding site. Binding to FIP200 alone in the absence of WIPI2 binding does not support autophagy induction (Dooley et al., 2014), and the nature of the interplay between FIP200 and WIPI2 binding to the ATG16L1 central region will be important to clarify. The Golgi-resident RAB33B also binds to ATG16L1 (Itoh et al., 2008), although the precise role of this interaction in autophagy is unclear. The RAB33B interaction was recently mapped structurally (Metje-Sprink et al., 2020), and the RAB33B binding site was found to terminate at ATG16L1 residue 210, just N-terminal to the first ordered residues in the W2IR. In principle, it seems possible that RAB33B, FIP200, and WIPI2 might be capable of binding simultaneously.

Orienting WIPI2d membrane in the edge-on geometry proposed on the basis of previous studies (Baskaran et al., 2012; Krick et al., 2012), the N-terminus of the W2IR projects in the direction opposite to the membrane (Fig. 9). This potentially positions the ATG16L1 coiled coil to project away from the PI(3)P-containing membrane to which WIPI2 is bound. One model is that ATG16L1 could conjugate LC3 to the nascent phagophore in *trans* whilst anchored to a PI(3)P-containing domain of the ER (Dooley et al., 2014). *In vitro*, however, it is possible for WIPI2 to efficiently stimulate LC3 lipidation PI(3)P containing membranes in *cis* (Fracchiolla et al., 2020). Given the possibility that the ATG16L1 coiled coil can pivot with respect to the W2IR, these structural data on their own do not rule *cis* or *trans* LC3 lipidation in or out. Additional structures of ATG16L1 as assembled with multiple regulators, in the context of the full ATG12– 5-16L1 complex, and in the context of membranes, will be required to answer this question. The high resolution structure presented here will be an important component for the interpretation of the larger scale, yet likely lower resolution, structures of assemblies yet to be solved.

**Figure 9:**
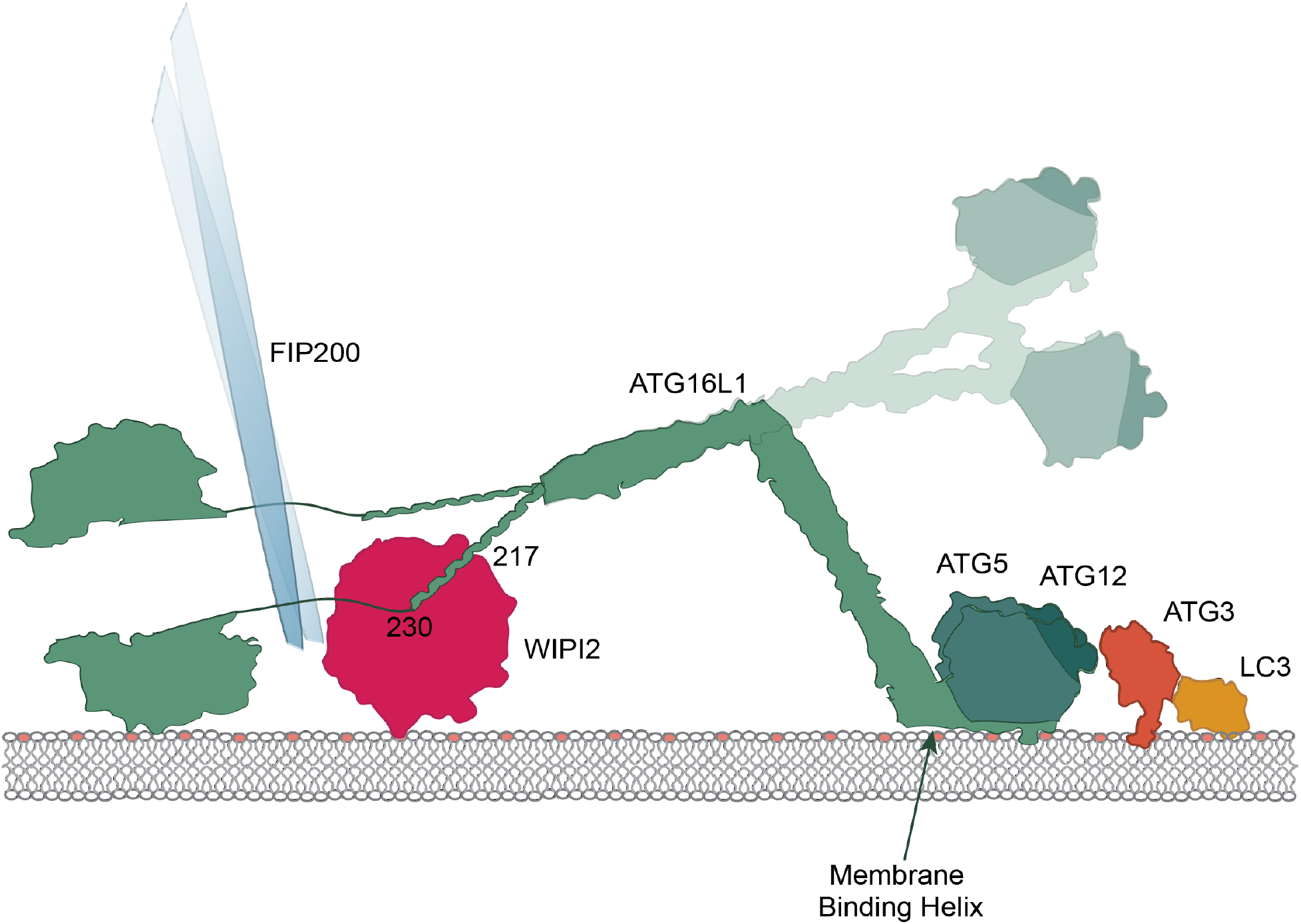
Model of WIPI2 recruitment of ATG16 to the membrane. Cartoon model of ATG16 positioning on the membrane while bound to WIPI2 and performing LC3 lipidation. Helix 1 membrane binding is labelled(Lystad et al., 2019), and a secondary upwards conformation is shown in faded colors. Rab33b binding is pictured.

## MATERIALS AND METHODS

### Plasmids

WIPI2d crystallography constructs and mutants were sub-cloned from a plasmid from a previous study(Fracchiolla et al., 2020) into the pCAG vector using restriction enzyme cloning. Mcherry constructs were cloned similarly with an N-terminal mcherry tag. All constructs had a C-terminal TEV cleavage site followed by TwinStrep tags.

### Protein expression and purification

Purification of WIPI2d constructs used for crystallization, pull-down assays, and GUV assays were expressed in HEK GnTi cells. Constructs were transfected to cells using polyethylenimine (Polysciences). After 60 h of expression, cells were harvested and lysed with lysis buffer (50 mM Hepes, pH 7.4, 1% Triton X-100, 300 mM NaCl, and 1 mM tris(2-carboxyethyl)phosphine [TCEP]) supplemented with EDTA-free protease inhibitors (Roche). The lysate was clarified by centrifugation (17,000 rpm for 1 h at 4°C) and incubated with StrepTactin Sepharose resin (IBA) for 2 h at 4°C, applied to a gravity column, and washed extensively with wash buffer (50 mM Hepes, pH 7.4, 300 mM NaCl, and 1mM TCEP). The protein complexes were eluted with wash buffer containing 10 mM desthiobiotin (Sigma) and treated with TEV protease at 4°C overnight. Cleaved protein was applied to a Superdex 200 column (16/60 prep grade) equilibrated with gel filtration buffer (25 mM Hepes, pH 7.4, 150 mM NaCl, and 1 mM TCEP). Peak fractions were collected, pooled, snap frozen in liquid nitrogen, and stored at −80°C. Purification of ATG12–5-16, PI3KC3-C1, ATG7, ATG3, and LC3 used for GUV assays were performed as previously described(Fracchiolla et al., 2020).

### Crystallization and Structural Determination

WIPI2d10-364Δ263-295: ATG16L1 (207-230) complex was formed overnight with 5X molar excess peptide (GenScript). Crystals of the complex were grown using hanging drop vapor diffusion method at 4°C. 1 μL of the protein complex (2 mg/mL) was mixed with 1 μL reservoir solution and 0.3 μL of a crystal seed stock. This was suspended over a 500 μL reservoir of 22% w/v PEG 3,350 (Hampton Research), 2% v/v Tacsimate pH 7.0 (Molecular Dimension), and 100mM Hepes pH 7.7. Crystals appeared within 2 days and were continued to grow for approximately a week. Crystals were cryoprotected in reservoir solution supplemented with 25% (v/v) glycerol. A native dataset was collected from a single crystal under cryogenic conditions (100°K) at a wavelength of 0.979Å using a Dectris PILATUS 6M/EIGER 16M detector (beamline BL12-2, SSRL). The data was indexed and integrated using LABELIT and XDS(Kabsch, 2010). Integrated reflections were scaled, merged, and truncated using AIMLESS and TRUNCATE, respectively. Initial phases were determined by molecular replacement with the program PHASER(McCoy et al., 2007) using KIHsv2 (PDB: 4EXV)(Baskaran et al., 2012) as a search model. ATG16L1 peptide was manually modeled into the structure according to the 2Fo-Fc and Fo-Fc electron density maps using Coot(Emsley et al., 2010). Iterative rounds of manual model building and refinement were performed using Coot(Emsley et al., 2010) and Phenix Refine(Afonine et al., 2012) respectively. Data collection and refinement statistics are listed in Supplementary Table 1. WIPI2 ATG16L1 interface was analyzed using PDBePISA(Krissinel and Henrick, 2007). All figures were generated with PyMol (http://www.pymol.org). The electrostatic surface was calculated using APBS(Baker et al., 2001) in PyMOL. Hydrophobic surface was generated using YBR script in PyMOL(Hagemans et al., 2015). WIPI1 and WIPI4 homology models were generated in SWISS-Model(Bertoni et al., 2017; Bienert et al., 2017; Studer et al., 2020; Studer et al., 2021; Waterhouse et al., 2018) using WIPI2d10-364Δ263-295 and WIPI3 (PDB: 6KLR) as templates, respectively.

### Coprecipitation Assay

10 μM purified WIPI2d was mixed with 20 μM of GST or GST-ATG16L1(207-230) and 10 μL Glutathione Sepharose 4B (GE Healthcare). The final buffer was 25mM HEPES pH 7.4, 150mM NaCl, 1mM TCEP. The final volume was 150 μL. The system was gently rocked at 4°C for 2 hours before washing the protein-bound resin three times. Loading dye was added to the beads and bands were visualized using SDS-PAGE gel after coomassie staining.

### GUV Assay

GUVs were prepared by hydrogel-assisted swelling as described previously (Chang et al, 2021). The reactions were set up in an eight-well observation chamber (Lab Tek) that pre-coated with 5 mg/ml β casein for 30 min. For E3 membrane recruitment and LC3 lipidation assay, a final concentration of 50 nM PI3KC3-C1 complex, 250 nM WIPI2d or mutant proteins, 50 nM E3-GFP complex, 100 nM ATG7, 100 nM ATG3, 500 nM mCherry-LC3B, 50 μM ATP, and 2 mM MnCl2 were used. For WIPI2d membrane binding assay, a final concentration of 50 nM PI3KC3-C1, 400 nM mCherry-WIPI2d or mutant proteins, 50 nM E3-GFP complex were used. A final volume of 120 μL mixture was made for all the reactions. 10 μL GUVs were added to initiate the reaction. After 5 min incubation, during which random views were picked for imaging, time-lapse images were acquired in multitracking mode on a Nikon A1 confocal microscope with a 63 × Plan Apochromat 1.4 NA objective. Three biological replicates were performed for each experimental condition. Identical laser power and gain settings were used during the course of all conditions.

For quantification of protein intensity on GUV membranes, the outline of individual vesicle was manually defined based on the membrane channel. The intensity threshold was calculated by the average intensities of pixels inside and outside of the bead and then intensity measurements of individual bead were obtained. Averages and standard deviations were calculated among the measured values per each condition and plotted in a bar graph. The data were analysed with GraphPad Prism 9 by using one-way ANOVA with Dunn’s multiple comparisons test.

### Starvation assay in MEFs

Wild-type SV40 immortalized MEFs were purchased from ATCC (CRL-2907) and cultured in DMEM (Corning) supplemented with 10% FBS. MEFs were plated on 35 mm glass bottom imaging dishes (MatTek) and on the following day transfected with 50 pmol ON-TARGETplus SMARTPool WIPI2 siRNA (Horizon) using RNAiMAX. After 24 hours, media was exchanged to fresh media and cells were transfected with 50 pmol WIPI2 siRNA and 0.75 μg of each Halo-WIPI2 construct using Lipofectamine 2000. 48 hours after Lipofectamine 2000 transfection, MEFs were starved in EBSS (Thermo Fisher) containing 100 nM bafilomycin A1 and 37.5 nM TMRDirect Halo Ligand (Promega). After 2 hours in EBSS, MEFs were fixed and permeabilized for 8 minutes at −20°C using ice-cold methanol. Cells were washed three times with PBS and blocked for 1 hour with 5% goat serum and 1% BSA in PBS. MEFs were then incubated with anti-LC3 primary antibody (Abcam) diluted in blocking solution for 1 hour at RT, washed three times with PBS, and incubated with anti-rabbit AlexaFluor488 secondary antibody for 1 hour at RT. After three washes with PBS and nuclear counterstaining with Hoechst (Thermo Fisher), MEFs were imaged in PBS on a Perkin Elmer spinning disk confocal setup with a Nikon Eclipse Ti inverted microscope, a Hamamatsu EMCCD 9100-50 camera, and an Apochromat 100x 1.49 NA oil immersion objective. Images were acquired as z stacks with a 200 nm step-size.

Z-stacks were assembled into maximal projections and channels were split using FIJI (NIH). Images from each condition across two biological replicates were used to train Ilastik to identify LC3 puncta. Images across three biological replicates (a unique passage of MEFs was considered a biological replicate) were used to train Ilastik to identify WIPI2 puncta after processing to normalize WIPI2 expression in FIJI. Training images were not used in subsequent data analysis. Fifteen images from each experiment for each condition were processed in batch mode by Ilastik to yield simple segmentation files. Using the WIPI2 channel, cell outlines were drawn by hand and saved as ROIs in FIJI. LC3 and WIPI2 puncta were counted within resulting ROIs using Analyze Particles in FIJI. For LC3 puncta, size was set 0-Infinity; for WIPI2 puncta, size was set 5-Infinity (square pixels). Results were tabulated in Microsoft Excel; graphing and statistical tests were performed using GraphPad Prism 9. Superplots were generated as discussed in Lord et al., 2020. One-way ANOVAs were performed on the averages for the biological replicates; Tukey’s multiple comparisons test was used post-hoc to compare WIPI2 point mutants to WT controls.

### Data Availability

Coordinates and structure factors have been deposited in the Protein Data Bank under accession code PDB 7MU2. Protocols will be deposited in protocols.io. Plasmids developed for this study will be deposited at Addgene.org. Other materials will be provided upon request to the corresponding author.

## ACKNOWLEDGEMENTS

We thank members of the Hurley and Holzbaur labs, Sascha Martens, Michael Lazarou and members of their laboratories, Dorotea Fracchiolla, and others in Aligning Science Across Parkinson’s Team mito911 for advice and discussions. We thank Clyde Smith and Lisa Dunn at SSRL beamline BL12-2 for assistance with data collection. The study is funded by the joint efforts of The Michael J. Fox Foundation for Parkinson’s Research (MJFF) and Aligning Science Across Parkinson’s (ASAP) initiative. MJFF administers the grant ASAP-000350 (to J.H.H. and E.H.) on behalf of ASAP and itself. C.A.B. was supported by the German Research Foundation (DFG; BO 5434/1-1). The research was also supported by National Institute of General Medical Sciences, NIH, R01 GM111730 (J.H.H.) and National Institute of Neurological Disease and Stroke, NIH, R00 NS109286 (A.K.H.S). Use of the Stanford Synchrotron Radiation Lightsource, SLAC National Accelerator Laboratory, is supported by the U.S. Department of Energy, Office of Science, Office of Basic Energy Sciences under Contract No. DE-AC02-76SF00515. The SSRL Structural Molecular Biology Program is supported by the DOE Office of Biological and Environmental Research, and by the National Institutes of Health, National Institute of General Medical Sciences (P30GM133894). The contents of this publication are solely the responsibility of the authors and do not necessarily represent the official views of NIGMS or NIH.

## CONTRIBUTIONS

L.S., C.C., C.A.B., A.S., C.Z.B., and T.G.F. performed research. L.S. and J.H. conceptualized research. E.H. and J.H. supervised research. L. S. and J. H. wrote the first draft of the manuscript. All authors commented upon the final draft of the manuscript.

---

**Supplementary Table 1.** Data collection and refinement statistics.

|  | WIPI2d |
| --- | --- |
| **Data Collection Statistics** |  |
| Wavelength | 0.9794Å |
| Resolution range | 38.29 - 1.85 (1.916 - 1.85) |
| Space group | I 2 |
| Unit cell | 117.8 49.1 120.1 90 95.9 90 |
| Total reflections | 126271 |
| Unique reflections | 55829 (1711) |
| Multiplicity | 3.3 |
| Completeness (%) | 97.6 (97.8) |
| Mean I/sigma(I) | 6.9 (2.7) |
| Wilson B-factor | 18 |
| R-merge | 0.05 (0.031) |
| R-meas | 0.098 |
| R-pim | 0.068 |
| CC1/2 | 0.996 |
| **Refinement statistics** |  |
| Reflections used in refinement | 55472 (5022) |
| Reflections used for R-free | 885 (85) |
| R-work | 0.1830 (0.2570) |
| R-free | 0.2188 (0.3225) |
| Number of non-hydrogen atoms | 5666 |
| macromolecules | 5041 |
| solvent | 625 |
| Protein residues | 658 |
| RMS(bonds) | 0.007 |
| RMS(angles) | 0.97 |
| Ramachandran favored (%) | 97.99 |
| Ramachandran allowed (%) | 2.01 |
| Ramachandran outliers (%) | 0 |
| Rotamer outliers (%) | 1.83 |
| Clashscore | 4.15 |
| Average B-factor | 25.37 |
| macromolecules | 24.26 |
| solvent | 34.32 |

\*Statistics for the highest-resolution shell are shown in parentheses.

**Supplementary Table 2:**
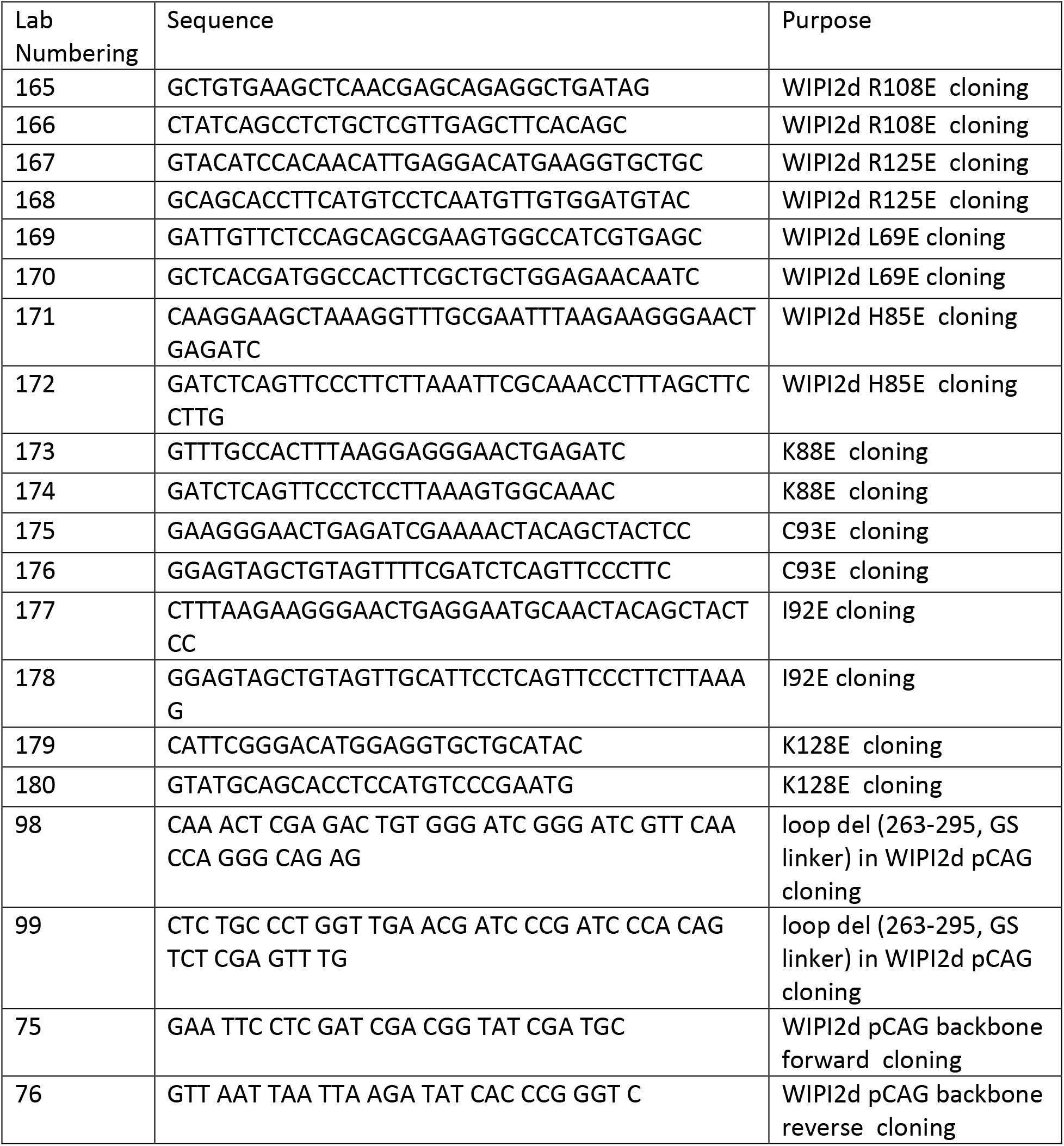
Oligos used for cloning

